# Restriction of dietary protein in rats increases progressive-ratio motivation for protein

**DOI:** 10.1101/2022.02.10.479961

**Authors:** Giulia Chiacchierini, Fabien Naneix, John Apergis-Schoute, James E. McCutcheon

## Abstract

Low-protein diets can impact food intake and appetite, but it is not known if motivation for food is changed. In the present study, we used an operant behavioral task – the progressive ratio test – to assess whether motivation for different foods was affected when rats were maintained on a protein-restricted diet (REST, 5% protein diet) compared to non-restricted control rats (CON, 18% protein). Rats were tested either with nutritionally-balanced pellets (18.7% protein, Experiment 1) or protein-rich pellets (35% protein, Experiment 2) as reinforcers. Protein restriction increased breakpoint for protein-rich pellets, relative to CON rats, whereas no difference in breakpoint for nutritionally-balanced pellets was observed between groups. When given free access to either nutritionally-balanced pellets or protein-rich pellets, REST and CON rats did not differ in their intake. We also tested whether a previous history of protein restriction might affect present motivation for different types of food, by assessing breakpoint of previously REST animals that were subsequently put on standard maintenance chow (protein-repleted rats, REPL, Experiment 2). REPL rats did not show increased breakpoint, relative to their initial encounter with protein-rich pellets while they were protein-restricted. This study demonstrates that restriction of dietary protein induces a selective increased motivation for protein-rich food, a behavior that disappears once rats are not in need of protein.

## 1. Introduction

The motivation to consume food strongly influences the amount of food consumed. In the context of maintaining homeostasis, increased motivation for food operates to restore energy or nutrient-specific depletion (Lutter & Nestler, 2009). In animal models, food restriction, for example, enhances motivation for highly caloric food (Jewett et al., 1995; Sharma et al., 2012). Similarly, in regards to sodium homeostasis, sodium depletion specifically increases operant responding for salt (Clark & Bernstein, 2006; Krieckhaus & Wolf, 1968; Quartermain et al., 1967) and enhances the motivational value of salt-associated cues (Robinson & Berridge, 2013).

The impact of dietary protein intake on cognitive functions is a subject of growing interest. In humans, maternal protein insufficiency causes offspring to have deficits in learning, memory and operant responding for a food reward (Grissom et al., 2014; Grissom & Reyes, 2013). Poorer cognitive functions in several domains (e.g. registration, attention, calculation, orientation, executive function) are reported in adults and older people on low-protein diets (Dickerson et al., 2020; Richard et al., 2018). In rodents, the importance of perinatal protein sufficiency for cognitive development has been demonstrated extensively (Almeida et al., 1996; Levitsky et al., 1975; McGaughy et al., 2014; Rushmore et al., 2021; Tonkiss et al., 1991a; Tonkiss & Galler, 1990; Tonkiss et al., 1991b). Notably, the effects of maternal protein malnutrition on spatial working memory and spatial learning are observed even trans-generationally (i.e. F_2_) (Abey et al., 2019). In adult rats, acute depletion of the essential amino acid tryptophan leads to impaired object recognition, increased anxiety and depression-related behavior (Jans et al., 2010). However, it is not known whether protein restriction involving a wider range of amino acids during adulthood causes cognitive deficits in rodents. In aging mice, protracted protein deficiency causes learning and memory deficits, which are reversed by essential amino acids administration (Sato et al., 2020). Overall, these studies indicate that cognitive impairments, especially in learning and memory, are strongly linked to protein deficiency, especially when the deficiency occurs during development. However, there is a lack of research investigating the consequences of protein restriction on motivation for food in rodents.

Our lab and others’ have recently demonstrated that adult rodents (> 2 months) maintained on a protein-restricted diet develop a strong preference for protein-containing food, relative to carbohydrate (Chiacchierini et al., 2021; Hill et al., 2019; Murphy et al., 2018; Naneix et al., 2020). Moreover, we recently showed that protein restriction during adulthood (postnatal day, P70-P82), but not during adolescence (P28-P42), increases dopamine release in the nucleus accumbens (Naneix et al., 2021). Furthermore, protein restriction in adulthood (>3 months) changes the response of ventral tegmental area neurons to the consumption of protein-containing food (Chiacchierini et al., 2021). What is not yet clear is whether this behavioral adaptation is also associated with changes in the motivation to obtain protein-rich food. Here, protein-restricted (REST) and control rats (CON) were trained to respond for pellets with differing protein content (nutritionally-balanced, 18% protein; protein-rich, 35% protein) and tested on a progressive ratio task in order to assess nutrient-specific changes in motivation. Additionally, we assessed whether a history of protein restriction affected motivation for protein-rich and nutritionally-balanced pellets when a nutritionally-balanced maintenance diet was restored.

## 2. Materials and Methods

### 2.1. Animals

Adult male Sprague Dawley rats were used for experiments (Experiment 1, n = 15; Experiment 2, n = 15. Charles River, weight range: 325-360 g; mean: 346 g at start of experiments). Rats were housed in pairs in individually ventilated cages (46.2 × 40.3 × 40.4 cm) with bedding material as recommended by NC3R guidelines. Temperature was 21 ± 2°C and humidity was 40-50%, with 12:12 h light/dark cycle (lights on at 07:00 am). Water and food were available ad libitum. Two rats were removed from the study because they did not show any instrumental learning during and after training (see section 2.6 for exclusion criteria). All experiments were covered by the Animals [Scientific Procedures] Act (1986) and carried out under the appropriate license authority (Project License: PFACC16E2).

### 2.2. Diets

All rats were initially maintained on standard laboratory chow (Teklad Global 18% Protein Rodent Diet, Envigo) (**Table 1**). A week after arrival, half of the rats were randomly assigned to the REST diet condition (Experiment 1, n=7; Experiment 2, n=7). For these rats, standard chow was switched to a modified AIN-93G diet containing 5% protein from casein (#D11092301, Research Diets; (Murphy et al., 2018) **Table 1**). Remaining rats were maintained under standard laboratory chow diet (CON; Experiment 1, n=8; Experiment 2, n=8). Behavioral testing started 1 week after diet manipulation.

**Table 1.** Maintenance diets used in the study. Macronutrient breakdown in control diet (Teklad Global; 18% protein) and protein-restricted diet (Research Diets #D11092301; modified AIN-93G; 5% protein).

|  | **Teklad Global** |  | **Research Diets D11092301** |  |
| --- | --- | --- | --- | --- |
|  | **(control, 18% protein)** |  | **(protein restricted, 5% protein)** |  |
|  | g (%) | kcal (%) | g (%) | kcal (%) |
| *Protein* | 18.6 | 24 | 5 | 4 |
| *Carbohydrate* | 44.2 | 58 | 76 | 74 |
| *Fat* | 6.2 | 18 | 10 | 22 |

### 2.3. Food reinforcers

Nutritionally-balanced pellets (#F0021, Bio-Serv) or protein-rich pellets (35% casein; #F07589, Bio-Serv) (**Table 2**) were used as reinforcers in Experiment 1 and Experiment 2, respectively.

**Table 2.** Reinforcers used in the study. Chemical composition of the food pellets used as reinforcers in Experiment 1 (#F0021) and in Experiment 2 (#F07589).

| **F0021 (nutritionally-balanced)** | **F07589 (casein-rich)** |
| --- | --- |
| 18.7% Protein | 35% Protein (Casein) |
| 59.1% Carbohydrate | 0.5% L-Methionine |
| 4.7% Fibre | 64.5% Fibre |
| 5.6% Fat |  |
| 6.5% Ash |  |
| < 10% Moisture |  |

### 2.4. Testing apparatus

Rats were tested in standard operant chambers (25 × 32 × 25.5 cm, Med Associates) placed inside sound attenuating chambers (1200 × 700 × 700 cm) with inbuilt ventilation fans. Each conditioning chamber was equipped with a house light located on the left wall while on the right wall there was a custom-designed pellet trough (6 × 6.5 × 2 cm; 3D printed using Open Scad 2015.03 and Ultimaker 2+) and a retractable lever (Med Associates), positioned either on the left or on the right of the pellet trough. The pellet trough was connected to a pellet dispenser (Med Associates) via a plastic tube. The position of the lever (right or left side) was counterbalanced between rats. The house light was turned on at the beginning of the session and turned off at the end of it. All behavioral tests were conducted during the light phase of the light/dark cycle, 5 days a week. Apparatus was controlled and data were recorded onto a PC using MED-PC IV software.

### 2.5. Magazine training

A week after diet manipulation started, rats were familiarized with the behavioral chamber and pellet delivery system through a magazine training session, in which 50 pellets were delivered into the pellet trough, at pseudo-random intervals (mean inter-pellet interval 40 ± 15 s), over a period of 45 minutes. The lever was retracted during the entire duration of the session.

### 2.6. Fixed ratio training

Twenty-four hours after magazine training, rats were trained on a fixed-ratio (FR) schedule of reinforcement, during which the lever was always extended. First, rats were trained to press the lever on a FR1 schedule, during which each response resulted in the delivery of one pellet. In subsequent sessions, rats progressed to FR2 (one pellet every two lever presses) and FR5 (one pellet every five lever presses) schedules. For each FR schedule, rats performed a daily session for 5 consecutive days. Reinforced responses were followed by a 5-second timeout period, during which lever presses did not result in additional pellet delivery but the number of lever presses was still recorded. Each FR session was terminated following 45 minutes or 100 pellets earned. Rats earning less than 5% of maximum rewards (i.e., 5 pellets) on at least three consecutive FR5 sessions were excluded from the study.

### 2.7. Progressive ratio testing

Twenty-four hours after the last training session, rats were tested under a progressive ratio 3 (PR3) schedule for 5 consecutive days. In this test, the number of lever presses required to earn the reinforcer increased progressively by 3 after each reinforcer was delivered, starting at 1 (i.e., 1, 4, 7, 10, etc.). Breakpoint was defined as the last ratio completed when no lever pressing occurred for more than 30 minutes or after 2 hours from the start of the session. Breakpoint is considered an index of motivation (Hodos, 1961).

### 2.8. Free access testing

Twenty-four hours after the last PR3 session, two daily free access tests were conducted. Rats were placed in the behavioral chambers with the house light on and the lever retracted. For 30 min they had free access to 15 g of pellets in the trough and their food consumption was measured.

### 2.9. Behavioral timeline

In Experiment 1, nutritionally-balanced pellets (see section 2.3) were used as reinforcers. Rats underwent the magazine training, fixed ratio training, progressive ratio testing and free access testing, as described in previous sections and in **Fig. 1A**.

**Figure 1.**
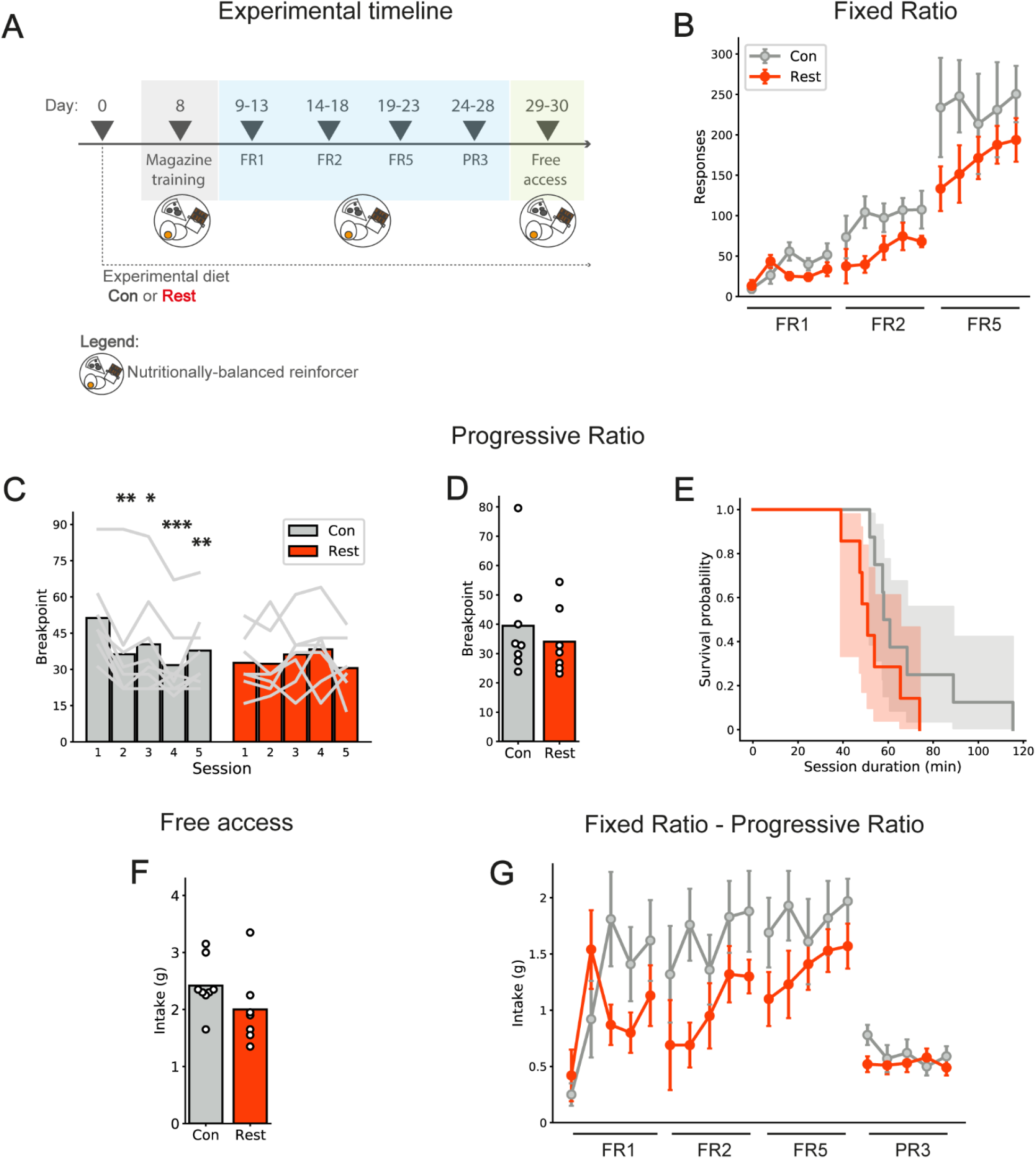
Protein restriction does not alter the motivation for food with nutritionally-balanced content. (A) Timeline of Experiment 1. (B) No difference between control (CON, grey, n = 8) and protein-restricted rats (REST, red, n = 7) in the number of responses made during fixed-ratio 1 (FR1), FR2 and FR5 sessions (mean ± SEM). (C) REST rats show constant breakpoint across five consecutive progressive ratio 3 (PR3) sessions. CON rats show a decrease in breakpoint across sessions. Bars show mean for each day and grey lines show data from individual rats. *, **, ***, p < 0.05, 0.01, 0.001 vs. Session 1 (Dunnett’s post hoc test). (D) No difference between CON and REST rats is observed in the average breakpoint across all days. Bars represent mean and circles represent individual values (rats). (E) Session duration is similar between CON and REST rats. Lines show survival curves for average session duration for all rats and shaded area is confidence interval. (F) CON and REST show similar intake of nutritionally-balanced pellets during free access. Bars show mean and circles represent individual values (rats). (G) Intake of reinforcers for fixed ratio and progressive ratio sessions (mean ± SEM).

Experiment 2 was performed with the same timeline of Experiment 1 but using protein-rich pellets as reinforcers. In addition, immediately after the last free access test, protein-restricted rats were placed back onto standard chow (protein repleted rats, REPL). After seven days on standard chow, both CON and REPL rats were tested on 5 daily progressive ratio sessions with protein-rich pellets, followed by 2 daily progressive ratio sessions with nutritionally-balanced pellets (**Fig. 1B**).

### 2.10. Statistical analysis

Number of responses, responses made during time out period and number of reinforcers delivered were recorded during fixed ratio and progressive ratio sessions. Breakpoints were recorded during progressive ratio sessions. The number of reinforcers left in the pellet trough was recorded at the end of each session, to discriminate between pellets delivered and pellets eaten. Intake of reinforcers in grams was calculated for each session by multiplying the number of pellets eaten by the weight of each pellet (45 mg). Statistical analysis was performed using GraphPad Prism 7 and SPSS 24. For the number of responses and the intake during fixed ratio sessions, three-way mixed ANOVA was used, with Diet as a between-subject variable, and Schedule and Session as within-subject variables. For breakpoints and intake during progressive ratio sessions, two-way mixed ANOVA was used with Diet as between-subject variable and Session as within-subject variable. Session duration was averaged across the five progressive ratio sessions for each animal, and compared between CON and REST rats with the Log-rank test. Pellet intake (free access tests), average breakpoints, post-reinforcement pause (i.e. time from reinforcer delivery to next lever press), and responses during timeout (progressive ratio tests) were averaged for each rat across sessions and compared between diet groups using unpaired t-tests. For summary data, CON and REST groups were obtained by pooling together animals from Experiment 1 and 2; for each animal, breakpoint was obtained by averaging all the progressive ratio sessions performed in a diet condition. Two-way mixed ANOVA was then used, with Diet as between-subject variable and Reinforcer type as within-subject variable. Significant effects and interactions were followed, if appropriate, with subsequent post hoc tests. All mixed ANOVAs were checked for sphericity of data using Mauchly’s Test and, if this was significant, the Huynh-Feldt corrected values were used. Assumptions of homogeneity of variance and normality were satisfied unless otherwise stated. Alpha was set at p < 0.05 and all significance tests were two-tailed. The number of animals was based on estimation from preliminary experiments.

### 2.11. Data and code availability

All data and custom analysis scripts are available at Zenodo and Github (DOIs: 10.5281/zenodo.5409201 and https://github.com/mccutcheonlab/PRPR.

## 3. Results

### 3.1. Experiment 1

#### 3.1.1. Protein restriction does not alter the motivation for nutritionally-balanced food

After magazine training, rats were trained to lever press for nutritionally-balanced pellets using FR1, FR2 and FR5 schedules. To ensure a similar level of training in all rats, each FR schedule was performed on five consecutive daily sessions (**Fig. 1A**). Throughout FR training, number of responses increased over the five sessions similarly in both groups (**Fig. 1B**). As such, three-way mixed ANOVA revealed a main effect of Session (F(4, 52) = 6.42, p < 0.0001), but no effect of Diet (F(1, 13) = 1.96, p = 0.184) or Schedule X Diet interaction (2, 13) = 1.44, p = 0.254). All other main effects and interactions were irrelevant to our hypothesis. The intake registered during sessions followed a similar pattern (**Fig. 1G**; Session, p < 0.001; Diet, p = 0.154; Schedule X Diet, p = 0.315). Following training on FR schedules and after 24 days on the protein-restricted diet for REST rats, rats were tested in five daily progressive ratio sessions, in which the number of lever presses required to earn the next reinforcer increased by three after each reinforcer delivery (PR3). We found that, across repeated PR3 sessions, CON and REST rats reached similar breakpoints. Moreover, breakpoint decreased across sessions in CON rats only (**Fig. 1C**). A two-way mixed ANOVA revealed a significant Diet X Session interaction (F(4, 52) = 6.32, p < 0.001), a main effect of Session (F(3.1, 40) = 3.58, p = 0.021) but no main effect of Diet (F(1, 13) = 0.47, p = 0.504). Subsequent multiple comparisons reported a significant decrease in breakpoint in CON rats across sessions (Dunnett’s post hoc tests vs. session 1: session 2, p = 0.004; session 3, p = 0.011; session 4, p < 0.001; session 5, p = 0.008) but not in REST rats (all Dunnett’s > 0.617). A similar trend was observed when the reinforcer intake during sessions was analyzed (**Fig 1G**; Diet X Session, p = 0.002; Session, p = 0.032; Diet, p = 0.487). Overall, when all five PR3 sessions were averaged together, the two diet groups did not differ in the motivation to obtain nutritionally-balanced reinforcers (t(13) = 0.69, p = 0.504) (**Fig. 1D**). We did not find any difference between groups in the number of responses made during the 5-second timeout (CON, 4.15 ± 2.84; REST, 4.94 ± 4.37; p = 0.680) and post-reinforcement pause (CON, 21.56 ± 15.77 s; REST, 18.89 ± 11.53 s; p = 0.719), indicating similar engagement in lever pressing behavior.

As the length of PR3 sessions also depended on animals’ engagement in lever pressing, we looked at the average duration of PR3 sessions as a further measure of motivation, and found that it was similar between CON and REST rats. The median survival rate for CON rats was 59 minutes and for REST rats it was 51 minutes (**Fig. 1E**). These survival curves were compared using a Log-rank test, which revealed no difference (p = 0.101), further supporting a similar motivation in the two diet groups to work for the pellets.

Following the five PR3 sessions and after 29 days on protein restriction for the experimental group, rats underwent two consecutive daily sessions of free access to the reinforcers (**Fig. 1F**). Pellet consumption across the two sessions was averaged for each rat. Unpaired t-test revealed no difference in the amount of reinforcers consumed between CON and REST rats (t(13) = 1.4, p = 0.177), indicating that protein restriction does not alter the intake of freely available nutritionally-balanced food. Even when the body weight of each animal was taken into account by calculating an intake index (grams consumed/grams of body weight), we did not observe any difference between diet groups (CON: 0.005 ± 0.0004, REST: 0.004 ± 0.0006; t(13) = 1.1, p = 0.302).

### 3.2. Experiment 2

#### 3.2.1. Protein restriction increases the motivation for protein-rich food

The second experiment was performed in a different cohort of rats, to investigate the effects of protein restriction on motivation specifically towards protein. Behavioral procedures were similar as in Experiment 1 but, instead of nutritionally-balanced pellets, protein-rich pellets were used (**Fig. 2A**). During training on FR schedules, REST rats displayed an increased number of lever presses, compared to CON rats (**Fig. 2B**). A three-way mixed ANOVA revealed a main effect of Diet (F(1, 13) = 6.61, p = 0.023) and a significant Schedule X Diet interaction (F(1, 13) = 5.76, p = 0.032). Analysis of reinforcer intake (**Fig. 2G**) revealed a main effect of Diet (p = 0.049), indicating that REST rats not only pressed more, but also consumed a greater amount of reinforcers during fixed-ratio sessions. No significant Schedule X Diet interaction was observed (p = 0.784).

**Figure 2.**
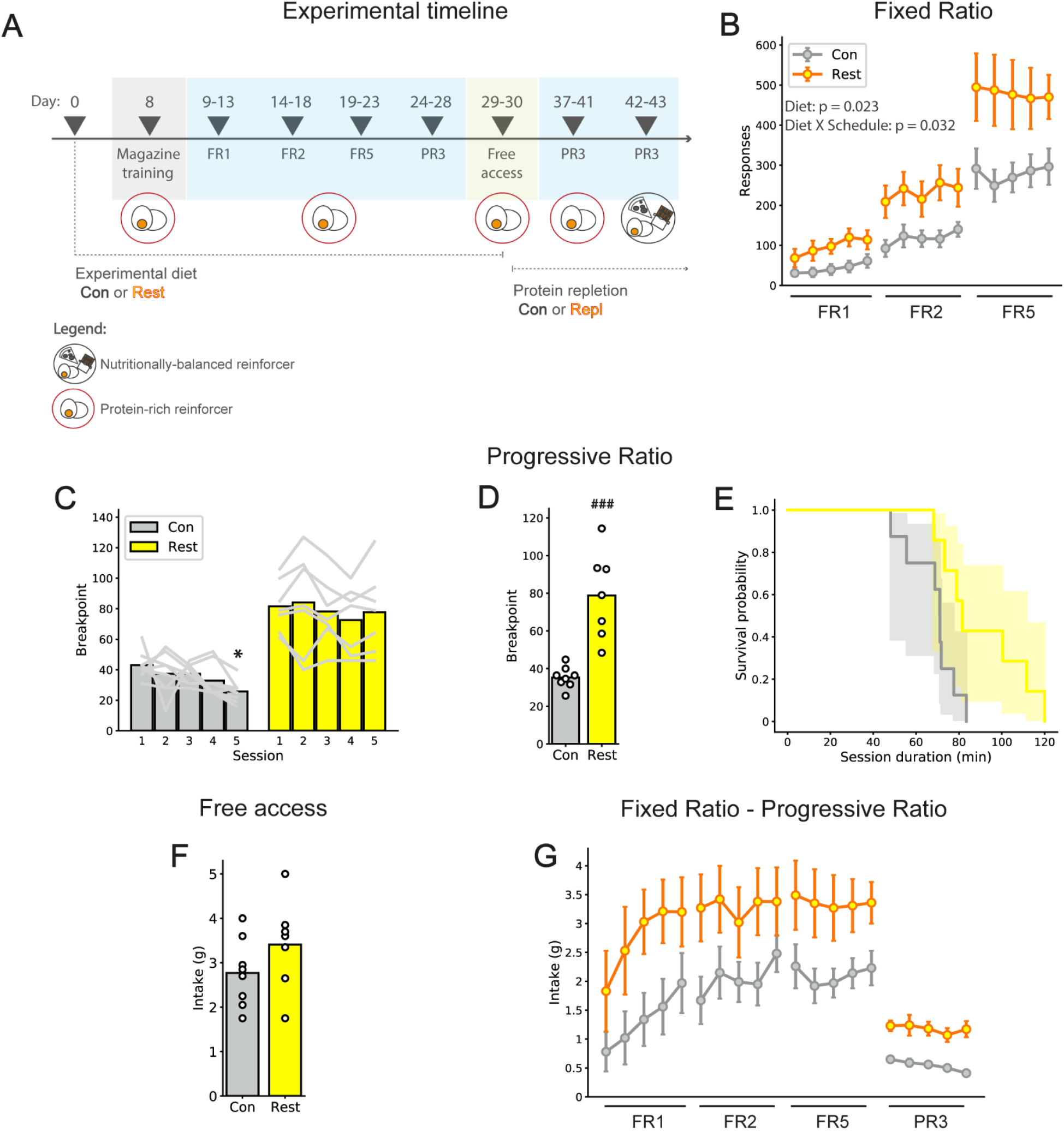
Protein restriction increases the motivation for protein-rich food. (A) Timeline of Experiment 2. (B) During FR sessions, protein-restricted (REST) rats show increased number of responses, compared to control (CON) rats (mean ± SEM). (C-D) During PR3 sessions, REST rats show elevated breakpoint, relative to CON rats. Bars show mean and grey lines (C) and circles (D) show data from individual rats. (*, p < 0.05 vs. Session 1, Dunnett’s post hoc test; ###, p < 0.001 vs. CON, unpaired t-test). (E) Progressive ratio session duration is longer in REST rats, compared to CON. Lines show survival curves for average session duration for all rats and shaded area is confidence interval. (F) During free access sessions, no difference between diet groups in intake is observed. Bars show mean and circles represent individual values (rats). (G) Intake of reinforcers for fixed ratio and progressive ratio sessions (mean ± SEM).

On progressive ratio (PR3) sessions, when the experimental group had been on the protein-restricted diet for 29 days, REST rats reached a higher breakpoint, relative to CON rats. (**Fig. 2C**). A two-way repeated measures ANOVA revealed a main effect of Diet (F(1, 13) = 26.9, p < 0.001) and of Session (F(2.34, 30.4) = 3.78, p = 0.028), but no significant interaction (F(4, 52) = 1.37, p = 0.257). The average breakpoint across sessions confirmed that REST rats were more motivated for protein than CON rats (t(13) = 5.19, p < 0.001) (**Fig. 2D**). REST rats also consumed more reinforcers during sessions, relative to CON rats (**Fig. 2G**; Diet, p = 0.0001). The increased motivation was also reflected in a higher survival rate of REST rats when the duration of progressive ratio sessions was analyzed (**Fig. 2E**). As such, the median survival rate of CON rats was 71 minutes, while for REST rats was 82 minutes. Comparison of survival curves revealed a significant difference (Log-rank test, p = 0.023).

Analysis of the number of responses made during timeout and the length of post reinforcement pauses identified no significant differences between diet groups (Timeout responses: CON, 6.8 ± 3.72; REST, 13.89 ± 11.51; p = 0.122; Post-reinforcement pause: CON, 16.17 ± 9.70 s; REST, 13.73 ± 2.76 s; p = 0.532). Interestingly, when rats were given free access to protein-rich pellets for 30 minutes (days 29 and 30 of protein-restricted diet), no difference in intake between diet groups was observed (unpaired t-test: t(13) = 1.40, p = 0.184) (**Fig. 2F**). Consistently, the intake index (g consumed/body weight) was similar between CON and REST rats (CON: 0.006 ± 0.0006, REST: 0.007 ± 0.0008; t(13) = 1.6, p = 0.137).

#### 3.2.2. Protein repletion abolishes the increased motivation for protein-rich food

Following the free access test, REST rats were switched back to regular maintenance chow (protein-repleted rats, REPL, **Fig. 2A**). After a week, both CON and REPL rats were tested again on PR3 schedule for protein-rich pellets, for five daily sessions. This allowed motivation for protein-rich food to be assessed in rats with a history of protein restriction, but after protein need state was abolished. We found that CON and REPL rats reached a similar breakpoint, which decreased across sessions (**Fig. 3**). As such, two-way repeated measures ANOVA revealed a main effect of Session (F(4, 52) = 15.3, p < 0.001), but no effect of Diet (F (1, 13) = 2.88, p = 0.114) and no interaction (F(4, 52) = 1.12, p = 0.359). Consistently, the duration of the session was now similar between CON and REPL rats (CON, 70 ± 20 min; REST, 74 ± 24 min; p = 0.722). Interestingly, an increase in breakpoint was observed in both diet groups (CON and REPL) when protein-rich reinforcers were replaced by nutritionally-balanced reinforcers (**Fig. 3, shaded columns**). A two-way repeated measures ANOVA revealed a main effect of Session (F(3.02, 39.2) = 13.6, p < 0.001), but no effect of Diet (F(1, 13) = 1.78, p = 0.205) and no significant interaction (F(6, 78) = 1.19, p = 0.321). Subsequent multiple comparisons indicated a progressive decrease in breakpoint, but the trend reverted to initial breakpoint value when nutritionally-balanced pellets were given (**Fig. 3**) (Dunnett’s post-hoc tests vs. Session 1: Session 2, p = 0.263; Session 3 to Session 5, all p_s_ < 0.006; Session 6 and 7, p_s_ > 0.405).

**Figure 3.**
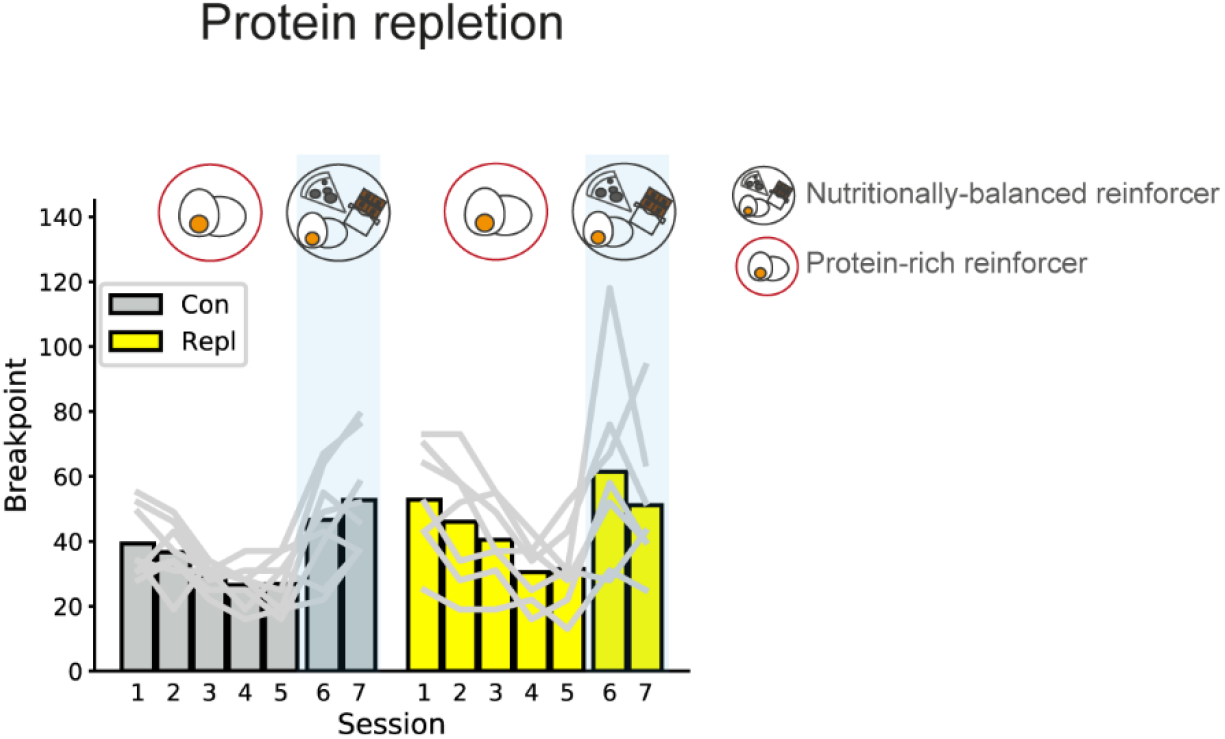
Protein repletion abolishes the increased motivation for protein food induced by protein need. Following experiment 2, all rats had access to regular maintenance chow for a week (controls, CON and protein-repleted rats, REPL). Both groups then underwent progressive ratio sessions for protein-rich (Session 1-5) and nutritionally-balanced reinforcers (Sessions 6 and 7). For protein-rich reinforcers, CON and REPL rats do not differ in breakpoint reached, which decreases across sessions similarly in both groups. However, when protein reinforcers are replaced by nutritionally-balanced reinforcers, both groups show a significant increase in breakpoint, but not different than each other. Bars show mean and grey lines are data from individual rats.

### 3.3 Comparison of progressive-ratio motivation for different reinforcers across all diet conditions

We next analyzed how breakpoint for nutritionally-balanced and protein-rich reinforcers changed according to the different dietary protein conditions: CON, REST and REPL. Protein status strongly and selectively influenced the motivation for food reinforcers, as shown by a main effect of Diet and a significant Diet X Reinforcer type interaction (**Fig. 4**, two-way mixed ANOVA: Diet, F(2, 27) = 4.56, p = 0.020; Diet X Reinforcer type, F(2, 27) = 31.0, p < 0.001; no main effect of Reinforcer type, p = 0.083). Further comparisons showed that only current protein restriction led to increased motivation for protein-rich pellets (Tukey’s post hoc tests: REST vs. CON, p < 0.001; REST vs. REPL, p < 0.001; CON vs. REPL, p = 0.610). Moreover, protein repletion induced an increase in the motivation for balanced reinforcers, relative to when rats were protein-restricted (Tukey’s post hoc tests: REST vs. REPL, p = 0.027; all other p_s_ > 0.2). Interestingly, for REPL rats, there was no significant difference in breakpoint between protein-rich and nutritionally-balanced reinforcers, suggesting that there is not a large difference in incentive value between them (Sidak’s post hoc test, p = 0.054, **Fig.4**), and indicating that, in REPL rats, protein-rich food has a similar incentive value as regular food, as observed in CON animals.

**Figure 4.**
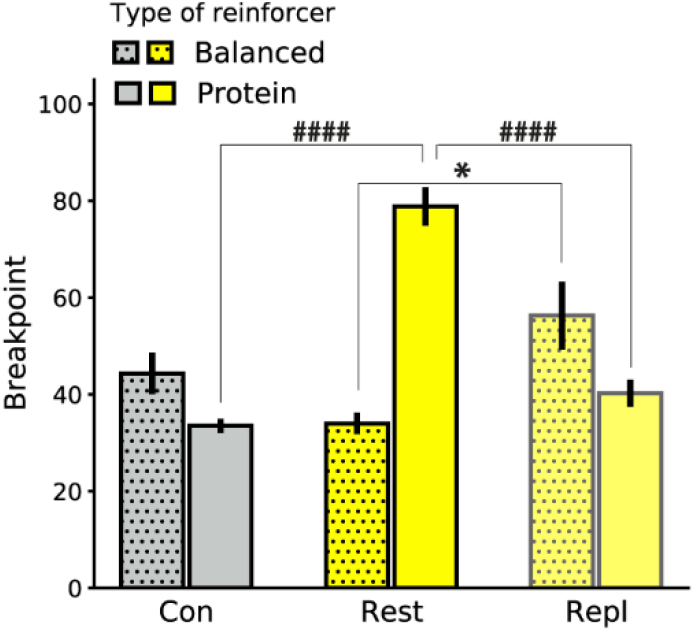
Current and previous protein status strongly and selectively influences motivation for food. Breakpoint for protein-rich reinforcers is elevated in protein-restricted (REST) rats only (yellow bar, center), compared to both control (CON; grey bar, left) and protein-repleted rats (REPL; pale yellow bar, right). Breakpoint for nutritionally-balanced reinforcers is elevated in REPL rats, relative to REST rats. No difference in breakpoint for nutritionally-balanced reinforcers is observed in CON vs. REST and CON vs REPL (dotted bars). For this summary analysis, CON and REST groups are obtained by pooling together animals from Experiment 1 and 2. For each rat, breakpoint is obtained by averaging all the progressive ratio sessions performed in a diet condition. Bars show mean ± SEM. *, p < 0.05 ####, p < 0.001

## 4. Discussion

The effect of protein restriction on progressive ratio motivation towards food has not yet been determined and this was the main goal of the current study. To summarize, we found that protein restriction increased the motivation to earn protein-rich food, but not food in general, indicating that protein restriction-induced changes in motivation are selective for protein-rich food. Moreover, feeding animals with a nutritionally-balanced diet after a period of protein restriction resulted in the abolition of elevated motivation for protein-rich food. Interestingly, despite there being an increased motivation for protein food, when food was freely available its intake was similar between REST and CON rats.

The fulfillment of homeostatic needs such as hunger, thirst or salt appetite is known to drive ingestion-related motivation (Berridge, 2004). As such, food- and water-restricted rodents show increased instrumental responding selectively for the relevant reinforcer (Eiselt et al., 2021; Olarte-Sánchez et al., 2015), demonstrating that depriving rodents of food and water leads to an increase in their incentive value. Similarly, sodium depleted animals are able to perform high-effort sodium-directed activity to restore sodium homeostasis (Quartermain et al., 1967; Schulkin, 1986). Our results suggest that rodents’ instrumental behavior also adapts to compensate for protein insufficiency.

In Experiment 1, nutritionally-balanced pellets were used as food reinforcers. During training under FR1, FR2 and FR5 schedules of reinforcement, CON and REST rats made a similar number of lever presses. Conversely, when protein-rich reinforcers were used (Experiment 2), REST rats made an increased number of lever presses already during training sessions. Thus, at a group level, increased lever pressing during FR sessions was associated with increased lever pressing during progressive ratio sessions. Although FR1 and FR2 are low effort schedules of reinforcement and are typically considered a measure of consummatory behavior rather than motivation (Arnold & Roberts, 1997), our data are consistent with other studies reporting a consistency between fixed ratio and progressive ratio measures of reward’s motivational properties (Fotio et al., 2021; Velázquez-Sánchez et al., 2014). As regards FR5, it has been proposed as a moderate-effort schedule measuring both intake and motivation (Vendruscolo et al., 2010), therefore the consistency found here between FR5 and PR3 is in support of this idea. Our results are also in line with previous findings of increased lever presses for protein-rich pellets during FR5 in golden hamsters fed a protein-free diet, relative to nutritionally complete-fed hamsters (DiBattista, 1999).

Stable performance on progressive ratio schedule is believed to require at least three sessions (Depoortere et al., 1993; Roberts et al., 1989). We performed five daily progressive ratio sessions and found that, while REST rats show a stable performance, CON rats showed a decrease in breakpoint across sessions, in both experiments. In Experiment 1, however, we observed elevated breakpoint in session 1 only and similarly reduced breakpoints on all the following sessions (**Fig. 1C**), suggesting that elevated breakpoint in the first session is driving the overall decrease. Instead, in Experiment 2, the decrease in breakpoint appears incremental across sessions (**Fig. 2C**), becoming statistically significant only on the last session. This decrease in the motivation to obtain protein-rich reinforcers is similar to what happens with calorie-free reinforcers (Beeler et al., 2012). Thus, it may be that, with experience, CON rats devalue protein-rich reinforcers due to the lack of other macronutrients in a similar way as rodents do when presented with reinforcers that do not provide nutritional benefit to the organism.

Intriguingly, we found increased breakpoint in both diet groups when working for the nutritionally-balanced reinforcers after a prior history of lever pressing for protein-rich reinforcers (**Fig. 3**, sessions 6 and 7). This might reflect a novelty effect of experiencing a new food option. In fact, when exposed to a novel food, rats take small food samples and, if the food is not associated with adverse body reactions, they increase the consumption (Mitchell, 1976). It is reasonable that our progressive ratio session lasts long enough to allow rats to sample and increase consumption of the novel nutritionally-balanced reinforcers.

When CON and REST rats were given nutritionally-balanced pellets in the food trough and were free to eat them for 30 minutes (Experiment 1), no difference between groups in total intake was observed. This result is in contrast with previous studies reporting increased food intake as a consequence of moderate protein restriction (Du et al., 2000; Morrison et al., 2012; White et al., 2000), which can be interpreted as a compensatory mechanism to make up for the lack of protein (Hill & Morrison, 2019; Simpson & Raubenheimer, 2005). Surprisingly, even when protein-rich reinforcers were used (Experiment 2), REST rats did not show increased intake during free access, despite an increased breakpoint, and consequently increased intake, during the progressive ratio task. This result might prove to be counterintuitive, especially in light of previous data from our lab (Chiacchierini et al., 2021; Murphy et al., 2018) and others (Chaumontet et al., 2018; Hill et al., 2019), showing an increased intake of protein-rich food, relative to carbohydrate, in REST rats when given the choice between the two nutrients. However, in the mentioned studies, protein and carbohydrate-rich food were simultaneously available, which may have resulted in a negative contrast effect (Mitchell & Flaherty, 1998) such as the value of carbohydrate, relative to protein, was decreased as a function of the comparison, leading to increased protein consumption. Conversely, in the present study, rats have free access to a single option (protein-rich food), therefore the lack of comparison with carbohydrate might have resulted in no increased intake in REST rats. In line with this idea, the lack of increased intake of protein in REST rats in the absence of a choice between nutrients has been previously reported by our lab during conditioning and forced-choice sessions, when only one nutrient-rich solution was available (Chiacchierini et al., 2021; Murphy et al., 2018). Another possibility to explain the discrepancy between instrumental responding and free access results in Experiment 2 is that protein restriction had the effect of making rats less sensitive to the cost associated with the protein reinforcers, thereby elevating the threshold at which rats can sustainably exert effort. In behavioral economics, this effect is known as “inelastic” demand (Hursh & Silberberg, 2008). It can finally be hypothesized that rats, over the 30-minute free access test, might have eaten until a maximum and stopped due to satiety, a mechanism that did not seem to be affected by protein restriction. This idea is supported by evidence that the average intake of rats during free access tests is much higher compared to intake during PR3 sessions (Average intake. Exp 1, Free access: CON 2.4 g; REST 2 g. Progressive ratio: CON 0.6 g; REST 0.5 g. Exp 2, Free access: CON 2.8 g; REST 3.4 g. Progressive ratio: CON 0.5 g; REST 1.2 g).

An important limitation of this study is the inclusion of only a single degree of protein restriction. It is notable, in fact, that different extents of protein restriction leads to different feeding behaviors in rodents, with moderately low-protein diets (between 5 and 10% protein) inducing hyperphagia (Morrison et al., 2007; White et al., 2000), while < 5% protein diets dramatically decrease food intake (Du et al., 2000; Wu et al., 2021; Zapata et al., 2019) - an effect that has been linked to reduced signaling in the hypothalamic hunger-related pathway (Wu et al., 2021). Therefore, further research should be undertaken to investigate the effects of different degrees of protein restriction on food-related motivation. A further drawback is the use of male rats only. In light of the different protein requirements in male and female rats at adulthood and during development (Leibowitz et al., 1991), and considering the importance of adequate protein intake during pregnancy in both human and rodents (Gould et al., 2018; Grissom & Reyes, 2013), future work should determine the impact of protein restriction on motivation in female rats.

We have demonstrated for the first time the direct consequences of protein restriction in adult rats on the motivation for different types of food. The next step would be to use this behavioral assay to gain insight into the central mechanisms underlying the increased motivation for protein-rich food induced by protein need state. Work from our group has recently demonstrated an elevated ventral tegmental area neural activity in REST rats consuming a protein-rich solution, relative to carbohydrate (Chiacchierini et al., 2021). In addition, others have reported increased c-Fos protein expression in the nucleus accumbens of REST rats after consuming a high-protein meal, compared to balanced-protein and low-protein meals (Chaumontet et al., 2018). Given the role of mesolimbic dopamine pathway in both the acute effects and learned properties of food rewards (Martel & Fantino, 1996; Tobler et al., 2005) and the involvement of this pathway in homeostatic feeding (Branch et al., 2013; Cone et al., 2014; Sharma et al., 2012), it is likely that changes in motivation induced by protein need are encoded by changes in mesolimbic dopamine. Accordingly, we also recently showed that protein restriction by itself induced specific changes of dopamine release in the nucleus accumbens, but not in the dorsal striatum (Naneix et al., 2021). Neuromodulators such as serotonin have also been shown to be influenced by dietary amino acids content (Markus, 2008). In light of the role of serotonin in the adaptive preference for protein food in flies (Vargas et al., 2010) and the involvement of serotonin transmission in the nucleus accumbens in the regulation of food-directed progressive ratio motivation (Pratt et al., 2012), it is possible that this neurotransmitter is involved in the motivation for protein observed in our REST rats. Finally, humoral signals such as fibroblast growth factor 21 (FGF21) have also been implicated in the response to dietary protein restriction. In particular, FGF21 is increased in both humans and rodents maintained on a protein-restricted diet (Laeger et al., 2014), and FGF21 signaling in the brain is necessary for the metabolic and behavioral adaptations to protein restriction (Hill et al., 2022; Hill et al., 2019). A possibility is that FGF21 interacts with brain pathways responsible for modulating adaptive effort-related behavior in response to protein restriction.

Over the past century, the study of macronutrients’ effect on body composition, weight control and on the development of obesity has highlighted the role of carbohydrate and fat in the diet. More recently, it has been proposed that exaggerated consumption of fat and sugar is a compensatory response to the reduction of absolute protein content in the diet, as animals would ingest food for reaching a protein target (Raubenheimer & Simpson, 2019; Simpson & Raubenheimer, 2005). Consistent with this, reports from rodent and human work have shown that protein intake is prioritized over fat and carbohydrate intake in the face of changes in diet composition, resulting in overconsumption of calories when diets are low in protein. In contrast to other studies demonstrating increased food intake in rodents on low-protein diets (Hill et al., 2019; Laeger et al., 2016), in the present work we did not observe an increase in nutritionally-balanced pellets in response to protein restriction. However, while in the above-mentioned studies food intake was registered daily, in the present work we measured consumption over 30 minutes of free access test.

Beyond increasing food consumption, reduced intake of dietary protein also affects metabolic responses, by improving glucose tolerance and slowing fat mass gain (Fontana et al., 2016), reducing the accumulation of white adipose tissue (Wanders et al., 2015), increasing insulin sensitivity (Solon-Biet et al., 2015), increasing energy expenditure (Laeger et al., 2014), reducing body weight gain (Hill et al., 2022). Additionally, the deleterious effects of inadequate protein diet on neurodevelopment and cognitive functions have been widely demonstrated (Gould et al., 2018; Grissom & Reyes, 2013). In light of this evidence, a better understanding of the impact of low protein diet on food-related behaviors and brain regions involved may help to address both health and disease conditions.

## Acknowledgements

The authors acknowledge the help and support from the staff of the Division of Biomedical Services, Preclinical Research Facility, University of Leicester, for technical support and the care of experimental animals as well as colleagues in the Department of Neuroscience, Psychology and Behaviour at the University of Leicester for their academic contribution. This work was funded by the Biotechnology and Biological Sciences Research Council [grant #BB/M007391/1 to J.E.M.], the European Commission [grant #GA 631404 to J.E.M.], The Leverhulme Trust [grant #RPG-2017-417 to J.E.M. and J.A-S.], and Tromsø Research Foundation [grant #19-SG-JMcC to J.E.M.).

## References

Abey, N. O., Ebuehi, O. A. T., & Imaga, N. O. A. (2019). Neurodevelopment and Cognitive Impairment in Parents and Progeny of Perinatal Dietary Protein Deficiency Models. Front Neurosci, 13, 826. doi:10.3389/fnins.2019.00826

Almeida, S. S., Tonkiss, J., & Galler, J. R. (1996). Prenatal protein malnutrition affects exploratory behavior of female rats in the elevated plus-maze test. Physiol Behav, 60(2), 675–680. doi:10.1016/s0031-9384(96)80047-3

Arnold, J. M., & Roberts, D. C. (1997). A critique of fixed and progressive ratio schedules used to examine the neural substrates of drug reinforcement. Pharmacol Biochem Behav, 57(3), 441–447. doi:10.1016/s0091-3057(96)00445-5

Beeler, J. A., McCutcheon, J. E., Cao, Z. F., Murakami, M., Alexander, E., Roitman, M. F., & Zhuang, X. (2012). Taste uncoupled from nutrition fails to sustain the reinforcing properties of food. Eur J Neurosci, 36(4), 2533–2546. doi:10.1111/j.1460-9568.2012.08167.x

Berridge, K. C. (2004). Motivation concepts in behavioral neuroscience. Physiol Behav, 81(2), 179–209. doi:10.1016/j.physbeh.2004.02.004

Branch, S. Y., Goertz, R. B., Sharpe, A. L., Pierce, J., Roy, S., Ko, D., … Beckstead, M. J. (2013). Food restriction increases glutamate receptor-mediated burst firing of dopamine neurons. J Neurosci, 33(34), 13861–13872. doi:10.1523/jneurosci.5099-12.2013

Chaumontet, C., Recio, I., Fromentin, G., Benoit, S., Piedcoq, J., Darcel, N., & Tomé, D. (2018). The Protein Status of Rats Affects the Rewarding Value of Meals Due to their Protein Content. J Nutr, 148(6), 989–998. doi:10.1093/jn/nxy060

Chiacchierini, G., Naneix, F., Peters, K. Z., Apergis-Schoute, J., Snoeren, E. M. S., & McCutcheon, J. E. (2021). Protein appetite drives macronutrient-related differences in ventral tegmental area neural activity. J Neurosci. doi:10.1523/jneurosci.3082-20.2021

Clark, J. J., & Bernstein, I. L. (2006). Sensitization of salt appetite is associated with increased “wanting” but not “liking” of a salt reward in the sodium-deplete rat. Behav Neurosci, 120(1), 206–210. doi:10.1037/0735-7044.120.1.206

Cone, J. J., McCutcheon, J. E., & Roitman, M. F. (2014). Ghrelin acts as an interface between physiological state and phasic dopamine signaling. J Neurosci, 34(14), 4905–4913. doi:10.1523/jneurosci.4404-13.2014

Depoortere, R. Y., Li, D. H., Lane, J. D., & Emmett-Oglesby, M. W. (1993). Parameters of self-administration of cocaine in rats under a progressive-ratio schedule. Pharmacol Biochem Behav, 45(3), 539–548. doi:10.1016/0091-3057(93)90503-l

DiBattista, D. (1999). Operant responding for dietary protein in the golden hamster (Mesocricetus auratus). Physiol Behav, 67(1), 95–98. doi:10.1016/s0031-9384(99)00043-8

Dickerson, F., Gennusa, J. V., 3rd, Stallings, C., Origoni, A., Katsafanas, E., Sweeney, K., … Yolken, R. (2020). Protein intake is associated with cognitive functioning in individuals with psychiatric disorders. Psychiatry Res, 284, 112700. doi:10.1016/j.psychres.2019.112700

Du, F., Higginbotham, D. A., & White, B. D. (2000). Food intake, energy balance and serum leptin concentrations in rats fed low-protein diets. J Nutr, 130(3), 514–521. doi:10.1093/jn/130.3.514

Eiselt, A.-K., Chen, S., Chen, J., Arnold, J., Kim, T., Pachitariu, M., & Sternson, S. M. (2021). Hunger or thirst state uncertainty is resolved by outcome evaluation in medial prefrontal cortex to guide decision-making.

Fontana, L., Cummings, N. E., Arriola Apelo, S. I., Neuman, J. C., Kasza, I., Schmidt, B. A., … Lamming, D. W. (2016). Decreased Consumption of Branched-Chain Amino Acids Improves Metabolic Health. Cell Rep, 16(2), 520–530. doi:10.1016/j.celrep.2016.05.092

Fotio, Y., Ciccocioppo, R., & Piomelli, D. (2021). N-acylethanolamine acid amidase (NAAA) inhibition decreases the motivation for alcohol in Marchigian Sardinian alcohol-preferring rats. Psychopharmacology (Berl), 238(1), 249–258. doi:10.1007/s00213-020-05678-7

Gould, J. M., Smith, P. J., Airey, C. J., Mort, E. J., Airey, L. E., Warricker, F. D. M., … Willaime-Morawek, S. (2018). Mouse maternal protein restriction during preimplantation alone permanently alters brain neuron proportion and adult short-term memory. Proc Natl Acad Sci U S A, 115(31), E7398–e7407. doi:10.1073/pnas.1721876115

Grissom, N., Bowman, N., & Reyes, T. M. (2014). Epigenetic programming of reward function in offspring: a role for maternal diet. Mamm Genome, 25(1-2), 41–48. doi:10.1007/s00335-013-9487-6

Grissom, N. M., & Reyes, T. M. (2013). Gestational overgrowth and undergrowth affect neurodevelopment: similarities and differences from behavior to epigenetics. Int J Dev Neurosci, 31(6), 406–414. doi:10.1016/j.ijdevneu.2012.11.006

Hill, C. M., Albarado, D. C., Coco, L. G., Spann, R. A., Khan, M. S., Qualls-Creekmore, E., … Morrison, C. D. (2022). FGF21 is required for protein restriction to extend lifespan and improve metabolic health in male mice. Nat Commun, 13(1), 1897. doi:10.1038/s41467-022-29499-8

Hill, C. M., Laeger, T., Dehner, M., Albarado, D. C., Clarke, B., Wanders, D., … Morrison, C. D. (2019). FGF21 Signals Protein Status to the Brain and Adaptively Regulates Food Choice and Metabolism. Cell Rep, 27(10), 2934–2947.e2933. doi:10.1016/j.celrep.2019.05.022

Hill, C. M., & Morrison, C. D. (2019). The Protein Leverage Hypothesis: A 2019 Update for Obesity. Obesity (Silver Spring), 27(8), 1221. doi:10.1002/oby.22568

Hodos, W. (1961). Progressive ratio as a measure of reward strength. Science, 134(3483), 943–944. doi:10.1126/science.134.3483.943

Hursh, S. R., & Silberberg, A. (2008). Economic demand and essential value. Psychol Rev, 115(1), 186–198. doi:10.1037/0033-295x.115.1.186

Jans, L. A., Korte-Bouws, G. A., Korte, S. M., & Blokland, A. (2010). The effects of acute tryptophan depletion on affective behaviour and cognition in Brown Norway and Sprague Dawley rats. J Psychopharmacol, 24(4), 605–614. doi:10.1177/0269881108099424

Jewett, D. C., Cleary, J., Levine, A. S., Schaal, D. W., & Thompson, T. (1995). Effects of neuropeptide Y, insulin, 2-deoxyglucose, and food deprivation on food-motivated behavior. Psychopharmacology (Berl), 120(3), 267–271. doi:10.1007/bf02311173

Krieckhaus, E. E., & Wolf, G. (1968). Acquisition of sodium by rats: interaction of innate mechanisms and latent learning. J Comp Physiol Psychol, 65(2), 197–201. doi:10.1037/h0025547

Laeger, T., Albarado, D. C., Burke, S. J., Trosclair, L., Hedgepeth, J. W., Berthoud, H. R., … Morrison, C. D. (2016). Metabolic Responses to Dietary Protein Restriction Require an Increase in FGF21 that Is Delayed by the Absence of GCN2. Cell Rep, 16(3), 707–716. doi:10.1016/j.celrep.2016.06.044

Laeger, T., Henagan, T. M., Albarado, D. C., Redman, L. M., Bray, G. A., Noland, R. C., … Morrison, C. D. (2014). FGF21 is an endocrine signal of protein restriction. J Clin Invest, 124(9), 3913–3922. doi:10.1172/jci74915

Leibowitz, S. F., Lucas, D. J., Leibowitz, K. L., & Jhanwar, Y. S. (1991). Developmental patterns of macronutrient intake in female and male rats from weaning to maturity. Physiol Behav, 50(6), 1167–1174. doi:10.1016/0031-9384(91)90578-c

Levitsky, D. A., Massaro, T. F., & Barnes, R. H. (1975). Maternal malnutrition and the neonatal environment. Fed Proc, 34(7), 1583–1586.

Lutter, M., & Nestler, E. J. (2009). Homeostatic and hedonic signals interact in the regulation of food intake. J Nutr, 139(3), 629–632. doi:10.3945/jn.108.097618

Markus, C. R. (2008). Dietary amino acids and brain serotonin function; implications for stress-related affective changes. Neuromolecular Med, 10(4), 247–258. doi:10.1007/s12017-008-8039-9

Martel, P., & Fantino, M. (1996). Mesolimbic dopaminergic system activity as a function of food reward: a microdialysis study. Pharmacol Biochem Behav, 53(1), 221–226. doi:10.1016/0091-3057(95)00187-5

McGaughy, J. A., Amaral, A. C., Rushmore, R. J., Mokler, D. J., Morgane, P. J., Rosene, D. L., & Galler, J. R. (2014). Prenatal malnutrition leads to deficits in attentional set shifting and decreases metabolic activity in prefrontal subregions that control executive function. Dev Neurosci, 36(6), 532–541. doi:10.1159/000366057

Mitchell, C., & Flaherty, C. (1998). Temporal dynamics of corticosterone elevation in successive negative contrast. Physiol Behav, 64(3), 287–292. doi:10.1016/s0031-9384(98)00072-9

Mitchell, D. (1976). Experiments on neophobia in wild and laboratory rats: a reevaluation. J Comp Physiol Psychol, 90(2), 190–197. doi:10.1037/h0077196

Morrison, C. D., Reed, S. D., & Henagan, T. M. (2012). Homeostatic regulation of protein intake: in search of a mechanism. Am J Physiol Regul Integr Comp Physiol, 302(8), R917–928. doi:10.1152/ajpregu.00609.2011

Morrison, C. D., Xi, X., White, C. L., Ye, J., & Martin, R. J. (2007). Amino acids inhibit Agrp gene expression via an mTOR-dependent mechanism. Am J Physiol Endocrinol Metab, 293(1), E165–171. doi:10.1152/ajpendo.00675.2006

Murphy, M., Peters, K. Z., Denton, B. S., Lee, K. A., Chadchankar, H., & McCutcheon, J. E. (2018). Restriction of dietary protein leads to conditioned protein preference and elevated palatability of protein-containing food in rats. Physiol Behav, 184, 235–241. doi:10.1016/j.physbeh.2017.12.011

Naneix, F., Peters, K. Z., & McCutcheon, J. E. (2020). Investigating the Effect of Physiological Need States on Palatability and Motivation Using Microstructural Analysis of Licking. Neuroscience, 447, 155–166. doi:10.1016/j.neuroscience.2019.10.036

Naneix, F., Peters, K. Z., Young, A. M. J., & McCutcheon, J. E. (2021). Age-dependent effects of protein restriction on dopamine release. Neuropsychopharmacology, 46(2), 394–403. doi:10.1038/s41386-020-0783-z

Olarte-Sánchez, C. M., Valencia-Torres, L., Cassaday, H. J., Bradshaw, C. M., & Szabadi, E. (2015). Quantitative analysis of performance on a progressive-ratio schedule: effects of reinforcer type, food deprivation and acute treatment with Δ?-tetrahydrocannabinol (THC). Behav Processes, 113, 122–131. doi:10.1016/j.beproc.2015.01.014

Pratt, W. E., Schall, M. A., & Choi, E. (2012). Selective serotonin receptor stimulation of the medial nucleus accumbens differentially affects appetitive motivation for food on a progressive ratio schedule of reinforcement. Neurosci Lett, 511(2), 84–88. doi:10.1016/j.neulet.2012.01.038

Quartermain, D., Miller, N. E., & Wolf, G. (1967). Role of experience in relationship between sodium deficiency and rate of bar pressing for salt. J Comp Physiol Psychol, 63(3), 417–420. doi:10.1037/h0024611

Raubenheimer, D., & Simpson, S. J. (2019). Protein Leverage: Theoretical Foundations and Ten Points of Clarification. Obesity (Silver Spring), 27(8), 1225–1238. doi:10.1002/oby.22531

Richard, E. L., Laughlin, G. A., Kritz-Silverstein, D., Reas, E. T., Barrett-Connor, E., & McEvoy, L. K. (2018). Dietary Patterns and Cognitive Function among Older Community-Dwelling Adults. Nutrients, 10(8). doi:10.3390/nu10081088

Roberts, D. C., Bennett, S. A., & Vickers, G. J. (1989). The estrous cycle affects cocaine self-administration on a progressive ratio schedule in rats. Psychopharmacology (Berl), 98(3), 408–411. doi:10.1007/bf00451696

Robinson, M. J., & Berridge, K. C. (2013). Instant transformation of learned repulsion into motivational “wanting”. Curr Biol, 23(4), 282–289. doi:10.1016/j.cub.2013.01.016

Rushmore, R. J., McGaughy, J. A., Amaral, A. C., Mokler, D. J., Morgane, P. J., Galler, J. R., & Rosene, D. L. (2021). The neural basis of attentional alterations in prenatally protein malnourished rats. Cereb Cortex, 31(1), 497–512. doi:10.1093/cercor/bhaa239

Sato, H., Tsukamoto-Yasui, M., Takado, Y., Kawasaki, N., Matsunaga, K., Ueno, S., … Kitamura, A. (2020). Protein Deficiency-Induced Behavioral Abnormalities and Neurotransmitter Loss in Aged Mice Are Ameliorated by Essential Amino Acids. Front Nutr, 7, 23. doi:10.3389/fnut.2020.00023

Schulkin, J. (1986). The evolution and expression of salt appetite. In The physiology of thirst and sodium appetite (pp. 491–496): Springer.

Sharma, S., Hryhorczuk, C., & Fulton, S. (2012). Progressive-ratio responding for palatable high-fat and high-sugar food in mice. J Vis Exp(63), e3754. doi:10.3791/3754

Simpson, S. J., & Raubenheimer, D. (2005). Obesity: the protein leverage hypothesis. Obes Rev, 6(2), 133–142. doi:10.1111/j.1467-789X.2005.00178.x

Solon-Biet, S. M., Mitchell, S. J., Coogan, S. C., Cogger, V. C., Gokarn, R., McMahon, A. C., … Le Couteur, D. G. (2015). Dietary Protein to Carbohydrate Ratio and Caloric Restriction: Comparing Metabolic Outcomes in Mice. Cell Rep, 11(10), 1529–1534. doi:10.1016/j.celrep.2015.05.007

Tobler, P. N., Fiorillo, C. D., & Schultz, W. (2005). Adaptive coding of reward value by dopamine neurons. Science, 307(5715), 1642–1645. doi:10.1126/science.1105370

Tonkiss, J., Foster, G. A., & Galler, J. R. (1991a). Prenatal protein malnutrition and hippocampal function: partial reinforcement extinction effect. Brain Res Bull, 27(6), 809–813. doi:10.1016/0361-9230(91)90213-4

Tonkiss, J., & Galler, J. R. (1990). Prenatal protein malnutrition and working memory performance in adult rats. Behav Brain Res, 40(2), 95–107. doi:10.1016/0166-4328(90)90002-v

Tonkiss, J., Galler, J. R., Shukitt-Hale, B., & Rocco, F. (1991b). Prenatal protein malnutrition impairs visual discrimination learning in adult rats. J Psychobiology, 19(3), 247–250.

Vargas, M. A., Luo, N., Yamaguchi, A., & Kapahi, P. (2010). A role for S6 kinase and serotonin in postmating dietary switch and balance of nutrients in D. melanogaster. Curr Biol, 20(11), 1006–1011. doi:10.1016/j.cub.2010.04.009

Velázquez-Sánchez, C., Ferragud, A., Moore, C. F., Everitt, B. J., Sabino, V., & Cottone, P. (2014). High trait impulsivity predicts food addiction-like behavior in the rat. Neuropsychopharmacology, 39(10), 2463–2472. doi:10.1038/npp.2014.98

Vendruscolo, L. F., Gueye, A. B., Darnaudéry, M., Ahmed, S. H., & Cador, M. (2010). Sugar overconsumption during adolescence selectively alters motivation and reward function in adult rats. PLoS One, 5(2), e9296. doi:10.1371/journal.pone.0009296

Wanders, D., Stone, K. P., Dille, K., Simon, J., Pierse, A., & Gettys, T. W. (2015). Metabolic responses to dietary leucine restriction involve remodeling of adipose tissue and enhanced hepatic insulin signaling. Biofactors, 41(6), 391–402. doi:10.1002/biof.1240

White, B. D., Porter, M. H., & Martin, R. J. (2000). Effects of age on the feeding response to moderately low dietary protein in rats. Physiol Behav, 68(5), 673–681. doi:10.1016/s0031-9384(99)00229-2

Wu, Y., Li, B., Li, L., Mitchell, S. E., Green, C. L., D’Agostino, G., … Speakman, J. R. (2021). Very-low-protein diets lead to reduced food intake and weight loss, linked to inhibition of hypothalamic mTOR signaling, in mice. Cell Metab, 33(6), 1264–1266. doi:10.1016/j.cmet.2021.04.016

Zapata, R. C., Singh, A., Pezeshki, A., Avirineni, B. S., Patra, S., & Chelikani, P. K. (2019). Low-Protein Diets with Fixed Carbohydrate Content Promote Hyperphagia and Sympathetically Mediated Increase in Energy Expenditure. Mol Nutr Food Res, 63(21), e1900088. doi:10.1002/mnfr.201900088

